# The economical lifestyle of CPR bacteria in groundwater allows little preference for environmental drivers

**DOI:** 10.1101/2021.07.28.454184

**Authors:** Narendrakumar M. Chaudhari, Will A. Overholt, Perla Abigail Figueroa-Gonzalez, Martin Taubert, Till L. V. Bornemann, Alexander J. Probst, Martin Hölzer, Manja Marz, Kirsten Küsel

**Affiliations:** Aquatic Geomicrobiology, Institute of Biodiversity, Friedrich Schiller University, Jena, Germany.; German Center for Integrative Biodiversity Research (iDiv) Halle-Jena-Leipzig, Leipzig, Germany.; Department for Chemistry, Environmental Microbiology and Biotechnology, Group for Aquatic Microbial Ecology (GAME), University Duisburg-Essen, Essen, Germany.; RNA Bioinformatics and High Throughput Analysis, Friedrich Schiller University, Jena, Germany.; European Virus Bioinformatics Center, Friedrich Schiller University, Jena, Germany.; FLI Leibniz Institute for Age Research, Jena, Germany.

**Keywords:** Candidate Phyla Radiation (CPR), *Cand.* Patescibacteria, Economic lifestyle, Metagenomics, Microbial ecology.

## Abstract

The highly diverse *Cand*. Patescibacteria are predicted to have minimal biosynthetic and metabolic pathways, which hinders understanding of how their populations differentiate to environmental drivers or host organisms. Their metabolic traits to cope with oxidative stress are largely unknown. Here, we utilized genome-resolved metagenomics to investigate the adaptive genome repertoire of Patescibacteria in oxic and anoxic groundwaters, and to infer putative host ranges.

Within six groundwater wells, *Cand*. Patescibacteria was the most dominant (up to 79%) super-phylum across 32 metagenomes obtained from sequential 0.2 and 0.1 µm filtration. Of the reconstructed 1275 metagenome-assembled genomes (MAGs), 291 high-quality MAGs were classified as *Cand*. Patescibacteria. *Cand*. Paceibacteria and *Cand*. Microgenomates were enriched exclusively in the 0.1 µm fractions, whereas candidate division ABY1 and *Cand*. Gracilibacteria were enriched in the 0.2 µm fractions. Patescibacteria enriched in the smaller 0.1 µm filter fractions had 22% smaller genomes, 13.4% lower replication measures, higher fraction of rod-shape determining proteins, and genomic features suggesting type IV pili mediated cell-cell attachments. Near-surface wells harbored Patescibacteria with higher replication rates than anoxic downstream wells characterized by longer water residence time. Except prevalence of superoxide dismutase genes in Patescibacteria MAGs enriched in oxic groundwaters (83%), no major metabolic or phylogenetic differences were observed based on oxygen concentrations. The most abundant Patescibacteria MAG in oxic groundwater encoded a nitrate transporter, nitrite reductase, and F-type ATPase, suggesting an alternative energy conservation mechanism. Patescibacteria consistently co-occurred with one another or with members of phyla Nanoarchaeota, Bacteroidota, Nitrospirota, and Omnitrophota. However, only 8% of MAGs showed highly significant one-to-one association, mostly with Omnitrophota. Genes coding for motility and transport functions in certain Patescibacteria were highly similar to genes from other phyla (Omnitrophota, Proteobacteria and Nanoarchaeota).

Other than genes to cope with oxidative stress, we found little genomic evidence for niche adaptation of Patescibacteria to oxic or anoxic groundwaters. Given that we could detect specific host preference only for a few MAGs, we propose that the majority of Patescibacteria can attach to multiple hosts just long enough to loot or exchange supplies with an economic lifestyle of little preference for geochemical conditions.

## Introduction

Metagenomic sequencing of diverse environments has enabled the recovery of genomic information from a vast majority of uncultivated microbial dark matter, significantly expanding the tree of life. *Cand.* Patescibacteria is a superphylum also known as Candidate Phyla Radiation (CPR) that constitutes a major portion of this expanded tree of life [1]. Patescibacteria, initially recovered from groundwater and aquatic sediments [2,3], are now shown to inhabit a broad range of surface and subsurface habitats, such as marine water, freshwater, freshwater beach sands [4] hydrothermal vents [5], cold-water geyser [6,7], plant rhizosphere [8], alpine permafrost [9], permafrost thaw ponds [10], and many more habitats [11] including the human oral cavity [12–14]. Nevertheless, they dominate the groundwater, where they comprise 20-70% of the total microbial community [15–18] along with thermokarst lakes [19] and hypersaline soda lake sediments [20].

Patescibacteria have small genomes characterized by predicted minimal biosynthetic and metabolic pathways, and are reported to have an anaerobic, fermentative lifestyle [21,22]. These traits may be responsible for their high abundance in nutrient-limited groundwater habitats, which are mainly anoxic. Interestingly, oxic surface soils are a major source of CPR bacteria inhabiting modern groundwater (stored within last 50 years) [23], as these organisms are easily mobilized into soil seepage water [17,24], but their metabolic traits to cope with oxidative stress are largely unknown. Divergent trends in the preference for several hydrochemical parameters or specific host preferences seem to result in the differentiation of CPR bacteria in groundwater [17]. Similarly, little species-level overlap of metagenome-assembled genomes (MAGs) across varying groundwater sites suggests that CPR communities differ based on specific environmental factors including host populations [18].

Most Patescibacteria cells are estimated to have ultra-small diameters ranging from 0.1 µm to 0.3 µm [11,15,21] with few exceptions like Saccharimonadia (candidate division TM7) that may be as large as 0.7 µm in diameter [25]. Small cell sizes of Patescibacteria accompanied by reduced genomes [3,21,22] suggest host-associated lifestyles. Indeed, specific studies on Patescibacteria isolates along with co-culture and microscopic analyses provided evidence of their symbiotic associations with other organisms e.g. with *Paramecium bursaria*, a ciliated protist in freshwater [26], or with Actinobacteria (*Actinomyces odontolyticus*, *Propionibacterium propionicus*, *Schaalia meyeri*) in the human oral cavity [12,27–29]. Similarly, CPR bacteria attach as episymbionts to putative bacterial hosts through pilin-like appendages in pristine groundwater [18].

In contrast, single cell genomic and biophysical observations from 46 globally distributed groundwater sites did not support the prevailing view that Patescibacteria are dominated by symbionts [11]. The authors suggest that their unusual genomic features and prevalent auxotrophies may be the result of ancestral, primitive energy metabolism that relies on fermentation. Additionally, genome streamlining in free-living prokaryotes in the open ocean is a known mechanism to reduce functional redundancy and conserve energy [30]. Minimizing energy expenditure and nutrient demands has constituted a selective advantage for *Prochlorococcus* in surface waters where nutrients are scarce at the expense of versatility and competitiveness in changing conditions [31], and the same could be true for CPR bacteria dominating oligotrophic subsurface waters. Thus, there is the need to disentangle which lineages of CPR bacteria are host-dependent and which are free-living, and how much variation in terms of lifestyle, metabolism and gene content exists between those which show a preference for certain geochemical conditions.

In this study, we took advantage of a well-studied modern groundwater system within the Hainich Critical Zone Exploratory (CZE) located in Thuringia, Germany [32], dominated by CPR bacteria, that exhibits large environmental gradients from oxic to anoxic conditions accompanied by different well-specific microbiomes [33]. Using 291 manually curated MAGs we aimed to identify the adaptive genomic repertoire of CPR bacteria. Sequential filtration was performed to gather clues about possible physical association of ultra-small Patescibacteria with larger sized host ranges. We also inferred putative hosts for Patescibacteria based on the co-occurrence patterns with other microorganisms within the transect, especially based on abundances of all the MAGs enriched in the 0.2 µm filter fractions.

## Results

### Patescibacteria represent more than 50% of all prokaryotes in Hainich groundwater

*Cand*. Patescibacteria dominated the groundwater community representing on average more than 50 ± 18% (range 23-79%) prokaryotes across 32 metagenomes obtained from groundwater of six wells that was sequentially filtered through 0.2 µm and 0.1 µm filters, based on the proportion of the quality-controlled metagenomic reads mapped to the 16S rRNA database (SILVA SSU rRNA Ref NR99) [34]. Three major classes within the phylum were detected: *Cand*. Parcubacteria/Paceibacteria (36.2 ± 17.1%, range 13-65.7%), *Cand*. Microgenomatia (7.2 ± 3.1%, range 2-12%) and candidate division ABY1 (3.2 ± 1.4%, range 1.1-5.1%). Patescibacteria were found to be highly abundant in both filter fractions. Their relative abundances were significantly higher (two-proportions z-test, p-value 1.16e-05) in the 0.1 µm filter fractions (67.6 ± 9.1%, range 54.1-78.5%) than in the 0.2 µm filter fractions (35.5 ± 8.9%, range 23.1-51.1%), (Figure 1).

**Figure 1:**
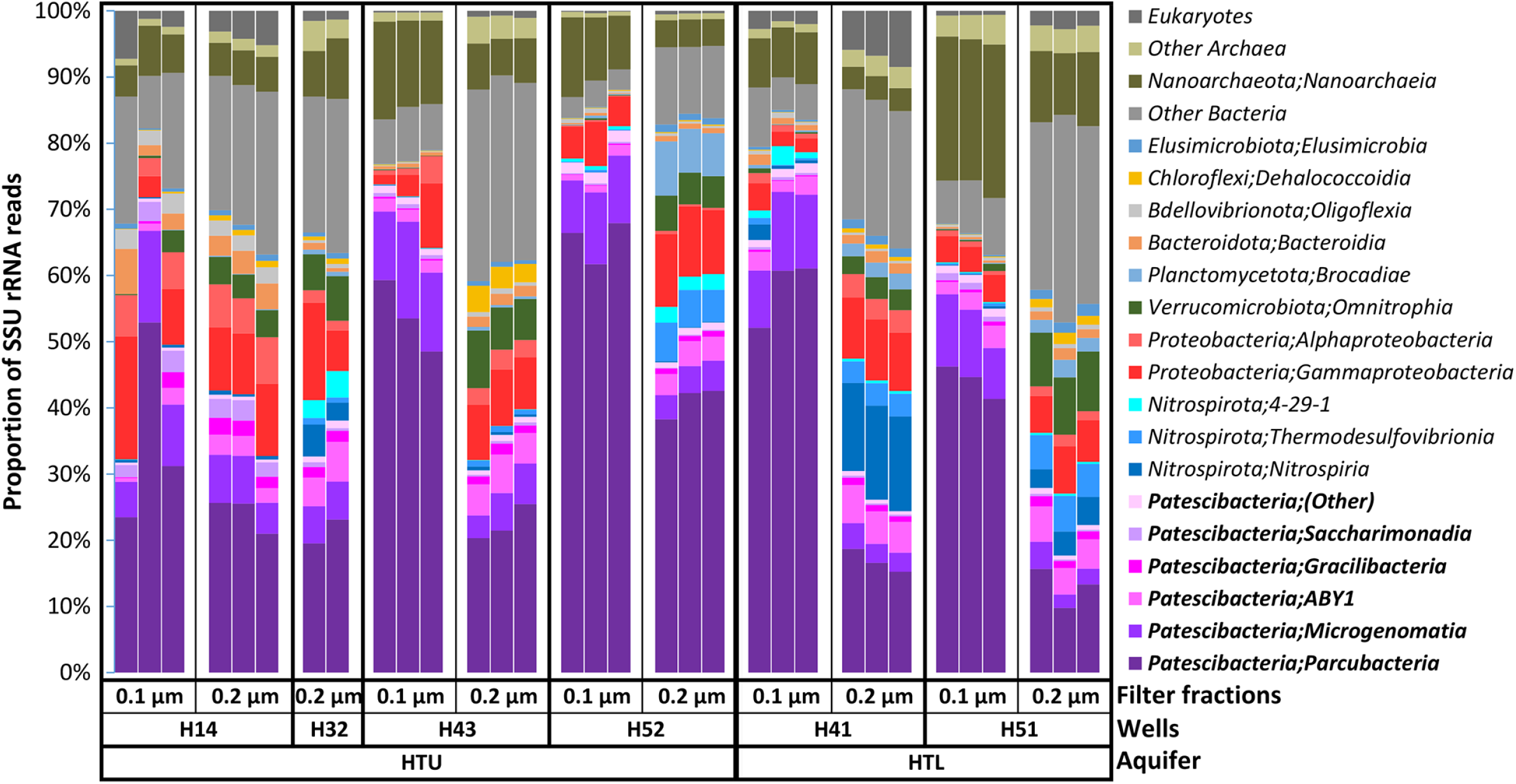
Community composition of the groundwater samples based on metagenomic reads mapped against the SILVA (SSU rRNA Ref NR99) database. Each column represents a metagenomic sample replicate for specified filter fractions from respective wells of the limestone-mudstone strata that host the multi-story upper aquifer assemblage (HTU; wells H14, H32, H43, H52) and the karstified main aquifer (HTL; wells H41, H51).

Within the detected Patescibacteria, site specific and filter size preferences were observed (Figure 2). The shallowest well at the top of the hillslope, H14, showed a relatively higher percentage of Saccharimondales compared to other wells. *Candidatus* Staskwiczbacteria showed preference for wells H14 and H43 (characterized by hypoxic/ anoxic environments with low nitrate), and *Candidatus* Wolfebacteria, UBA9983, and *Candidatus* Liptonbacteria for well H52 (characterized by anoxic environment and longest water residence time). *Candidatus* Magasanikbacteria and UBA9983 showed preference for 0.2 µm filter fractions of all the wells, whereas *Candidatus* Woesebacteria was enriched in all the 0.1 µm filter fractions.

**Figure 2:**
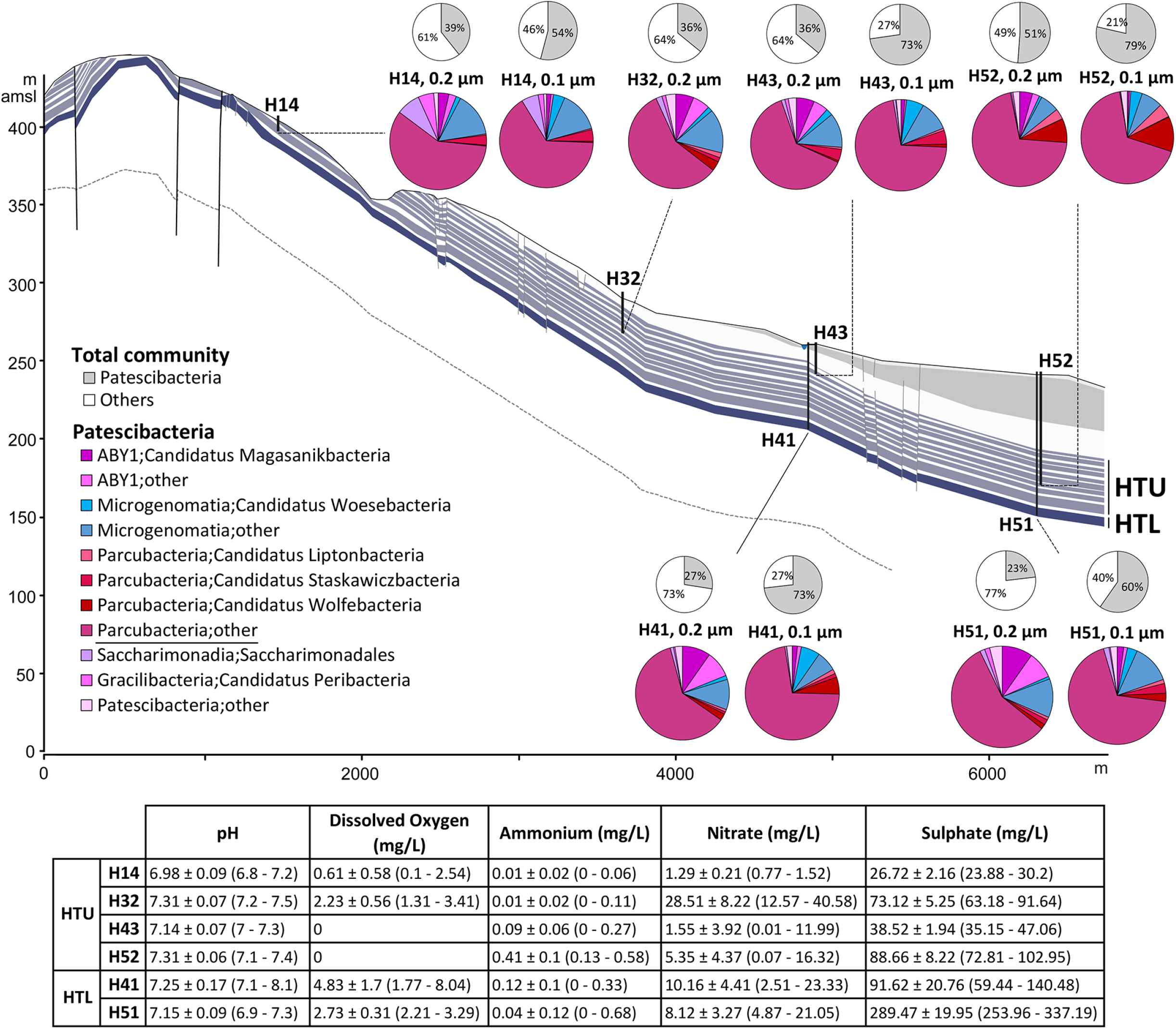
Community composition showing taxonomic preferences of Patescibacteria in wells and filter fractions across the Hainich transect. The cross section of the studied groundwater transect (from Kohlhepp *et al.*, 2017 [44], modified) shows the karstified main aquifer [HTL; (wells studied: H41, H51)] that is characterized by higher surface-connection to preferential recharge areas and the hanging thin- bedded alternating limestone-mudstone strata that host the multi-story upper aquifer assemblage (HTU; wells studied H14, H32, H43, H52). Height above mean sea level (amsl), in meters, is shown along the y-axis and length of hillslope is shown in meters along the x-axis. The colored pie charts show percentages of taxa within Patescibacteria at order level. The underlined taxon, *Parcubacteria;other* (all Parcubacteria other than the mentioned Parcubacteria orders merged together) was most abundant among Patescibacteria in all the filter fractions of all the wells. The grey pie charts show the relative percentage of Patescibacteria in the total community. The table includes levels of various hydrochemical parameters of the studied wells, including the dissolved oxygen, measured during July 2014 - April 2017 [33].

### Dominance of Patescibacteria in Hainich groundwater communities enabled recovery of hundreds of high quality MAGs

Metagenomic assembly and binning of all individual groundwater samples (n = 32) yielded a total of 1275 non-redundant manually refined MAGs from various bacterial and archaeal species. Among these MAGs, 584 MAGs were classified as *Cand*. Patescibacteria by GTDB-Tk and 291 of them were classified as CPR with high confidence score by a random forest classifier within Anvi’o v6.1 [35,36], trained with a set of CPR specific single copy genes extracted from previously published CPR genomes [15,37] (Additional file 1). Most of these 291 MAGs belonged to the classes: *Cand*. Paceibacteria (163 MAGs) followed by candidate division ABY1 (49 MAGs), and *Cand*. Microgenomatia (46 MAGs) (Figure 3, A). The details about all the Patescibacteria MAGs are provided in Additional file 2. The phylogenetic tree constructed from the multiple alignment of 68 core protein sequences confirmed the taxonomic placement of Patescibacteria MAGs (Figure 3, B).

**Figure 3:**
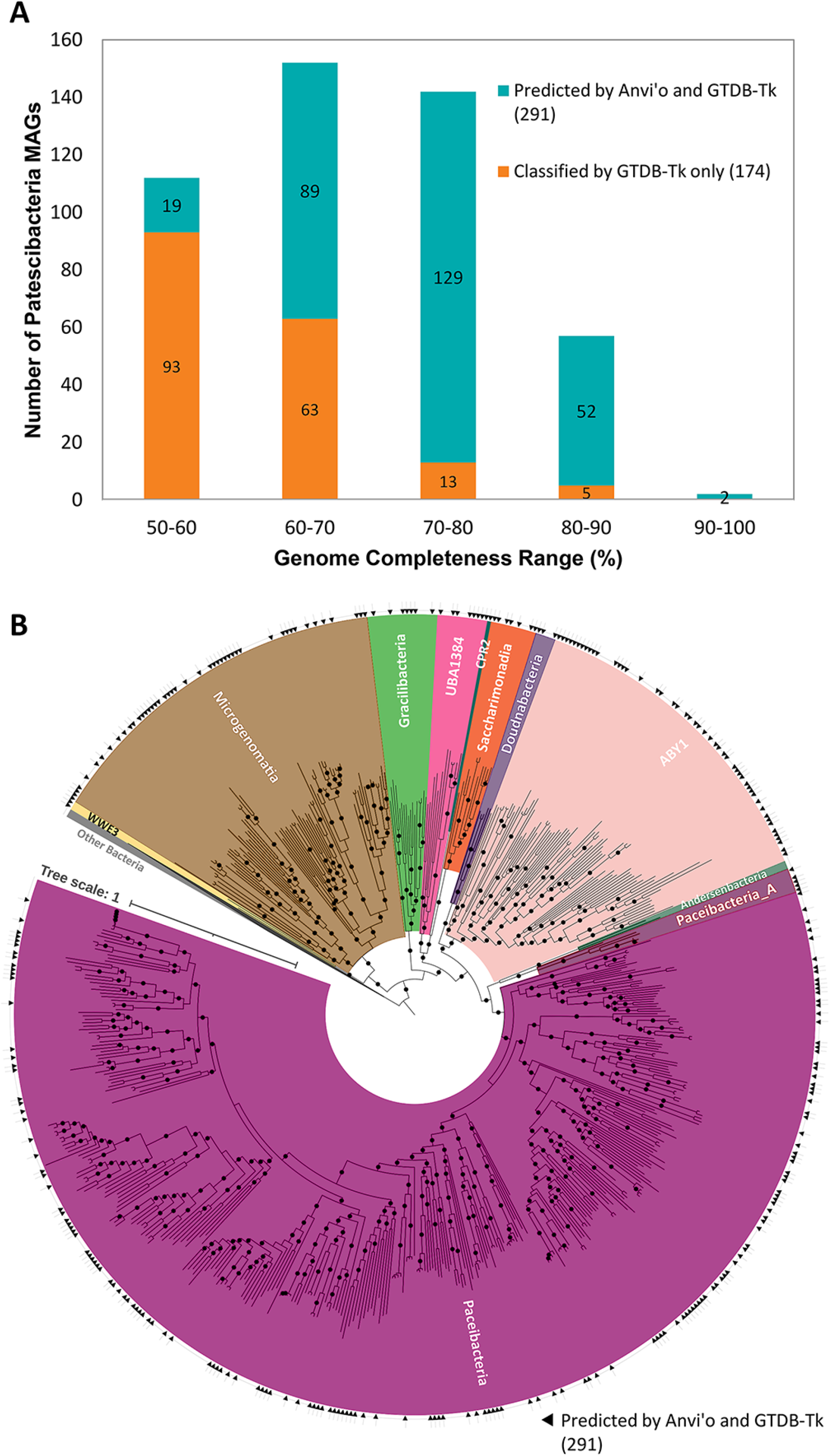
Phylogenetic placement of Patescibacteria MAGs after binning and refinement. **A.** Genome completeness distribution of the MAGs classified as Patescibacteria by GTDB-Tk alone (174, orange-colored bars), and by both GTDB-Tk and Anvi’o (291, teal-colored bars). **B.** Phylogenetic tree based on 68 core proteins from all bacterial MAGs (1087) using Maximum Likelihood in FastTree2 with 1000 bootstrap replications. Bacterial taxa other than Patescibacteria were collapsed together and only Patescibacteria are colored as per their taxonomic assignments from GTDB-Tk. The bootstrap values of 0.9 and above are indicated by filled circles. The phylogenetic tree is supplied as Additional file 3.

### Differences in the genome sizes of Patescibacteria based on cell size enrichment

We identified 110 Patescibacteria MAGs enriched in the 0.1 µm filter fractions based on their average *rpoB* gene-count-normalized coverage (See Methods) being 5-fold higher than in the 0.2 µm filter fractions. Of these, 82 MAGs were further classified as *Cand*. Paceibacteria, and 23 as *Cand*. Microgenomatia. Both classes were absent in the MAGs enriched in 0.2 µm filter fractions. Similarly, 33 Patescibacteria MAGs were enriched 5-fold more in the 0.2 µm filter fractions, with 22 of those belonging to the candidate division ABY1, and 5 to *Cand*. Gracilibacteria. Again, none of the genomes classified in these two classes were enriched in the 0.1 µm filter fractions.

The average genome size of all Patescibacteria MAGs enriched in the 0.1 µm filter fractions (688.7 ± 139.4 kb) was significantly smaller (Dunn’s test, p = 1.02e-06) than that of the Patescibacteria MAGs enriched in the 0.2 µm filter fractions (883.1 ± 204.3 kb), (Figure 4, A). There was no significant difference in the genome completeness and contamination values between the two groups.

**Figure 4:**
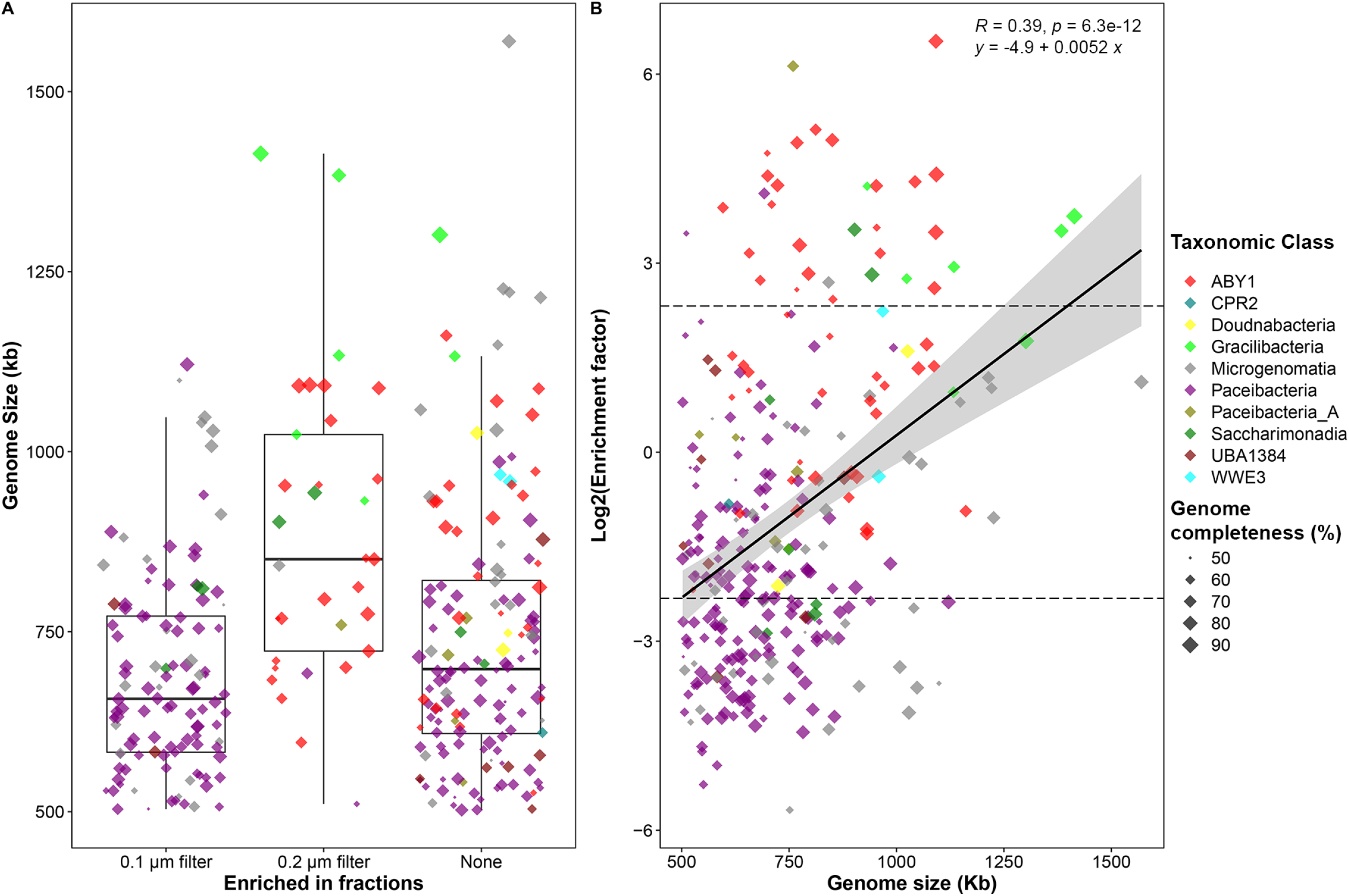
Distribution of genome sizes of Patescibacteria MAGs enriched in 0.1 µm and 0.2 µm filter fractions. **A**. For all 291 high-quality Patescibacteria MAGs, the ratio of average normalized genome coverage in 0.1 µm filter fractions to 0.2 µm filter fractions from metagenomes was used to form three groups: ‘0.1 µm filter’ - MAGs where this ratio was at least 5, ‘0.2 µm filter’ - MAGs where this ratio was ⅕ or less, and ‘None’ - MAGs other than first two groups. The mean genome sizes were significantly different (Kruskal-Wallis rank sum test, p = 2.24e-06). Pairwise Dunn’s test showed the genome sizes were significantly different between ‘0.1 µm filter’ and ‘0.2 µm filter’ (fdr adjusted p = 1.02e-06), and between ‘0.2 µm filter’ and ‘None’ (fdr adjusted p = 9.08e-05). **B.** The scatter plot shows the distribution of log2 filter enrichment factors (the ratio of average normalized genome coverage in 0.2 µm filter fractions to 0.1 µm filter fractions from metagenomes) of Patescibacteria MAGs, as the function of their genome sizes. The dashed lines indicate the cut-off value of 5 and ⅕ for filter enrichment factors on the y-axis.

When we analyzed the gene compositions of the two sets of Patescibacteria genomes, the genes encoding type-IV pilus assembly proteins (PilC, PilM, PilO) were significantly overrepresented (two-proportions z-test, p = 1.4e-04) in Patescibacteria enriched in the 0.1 µm filter fractions (∼88% of these genomes) as compared to those from the 0.2 µm filter fractions (∼64% of these genomes). Similarly, genes encoding cell division proteins FtsW and FtsI were present in 93% and 36% of the Patescibacteria MAGs enriched in 0.1 µm filter fractions, respectively. In comparison, the same genes were present in only 70% and 3% MAGs enriched in the 0.2 µm filter fractions (two-proportions z-test, p = 6.2e-04 and 4.7e-04). The gene encoding for the rod- shape determining protein (MreB) was also more likely to be found in Patescibacteria MAGs enriched in the 0.1 µm filter fraction (95% in the 0.1 µm-enriched vs 75% in the 0.2 µm- enriched, two-proportions z-test, p = 1.8e-03). Additionally, genes involved in colanic acid biosynthesis (*wcaH* and *wcaF*) were uniquely present in ∼10% of the Patescibacteria enriched in the 0.1 µm filter fractions.

Conversely, the L-lactate dehydrogenase gene was detected in 12% of the MAGs enriched in the 0.2 µm filter fractions and was entirely absent in the 0.1 µm-enriched MAGs. A similar pattern was found for the tryptophan synthase genes, *trpA* and *trpB*, which were detected in 15% and 18% of the MAGs enriched in the 0.2 µm filter fractions, but absent in Patescibacteria MAGs enriched in the 0.1 µm filter fractions.

### Growth dynamics of Patescibacteria using *in situ* measure of replication

Patescibacteria MAGs had comparatively higher estimated growth measures (GRiD values) in the near surface wells of the groundwater transect (wells H14 and H32), in comparison to the downstream wells (Figure 5, A). Specifically, these Patescibacteria showed significantly higher GRiD values at well H14 as compared to the downstream wells H41 and H43, and significantly higher GRiD values at well H32 as compared to all other wells present downstream. Notably, the wells with highest mean GRiD values for Patescibacteria were also the wells with lowest number of Patescibacteria MAGs. (Additional file 4, Figure S1).

**Figure 5:**
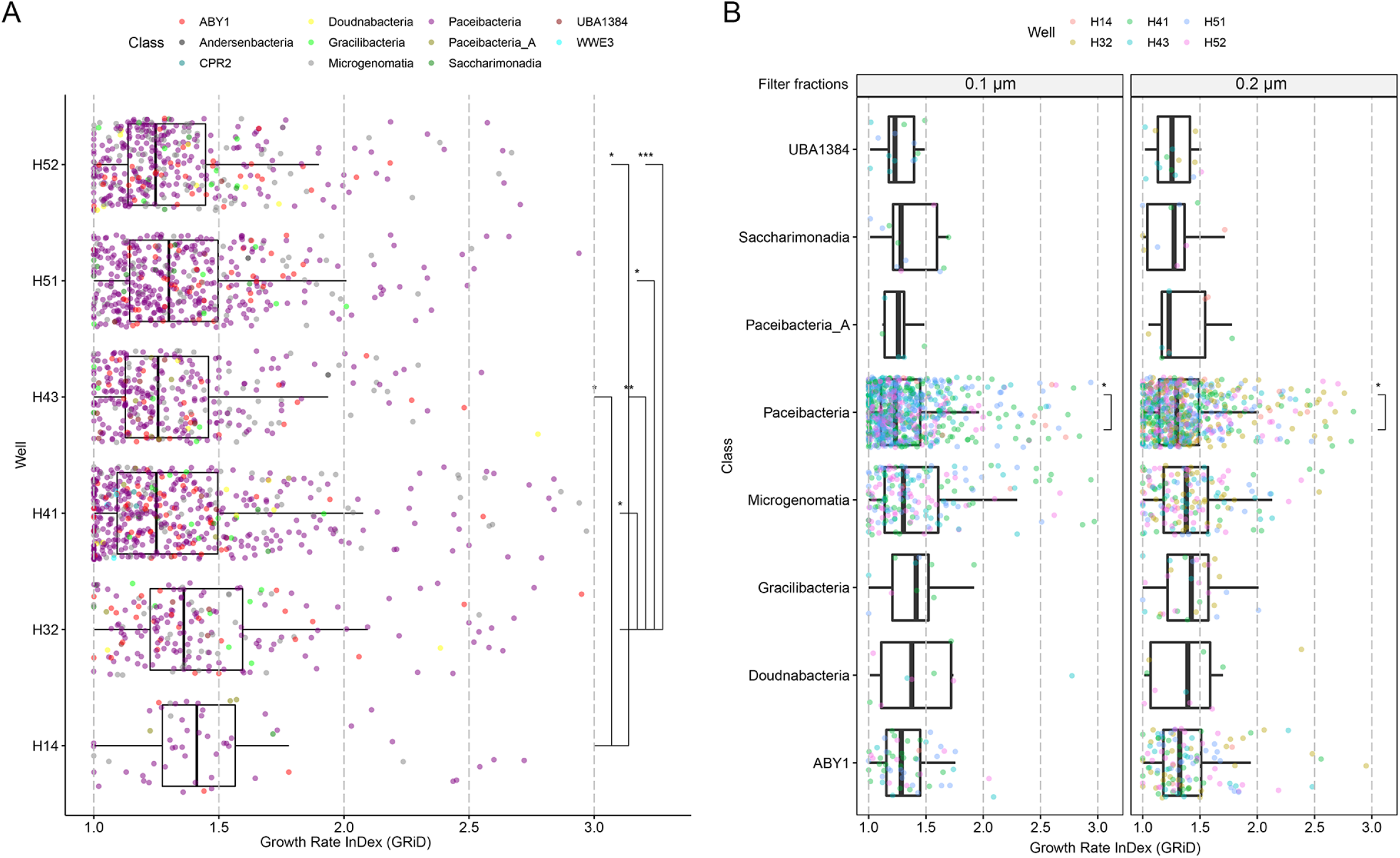
The estimated growth rate index (GRiD) distribution of Patescibacteria MAGs across the metagenomes. **A.** Well-wise GRiD distribution of all Patescibacteria. **B.** GRiD distribution of classes of Patescibacteria in 0.1 µm and 0.2 µm filter fractions. The statistical significance was calculated by using the t_test function with FDR correction in R package *rstatix* [78].

The GRiD values were significantly higher (Welch Two Sample t-test, p = 8.73e-07) in Patescibacteria MAGs enriched in 0.2 µm filter fractions (1.40 ± 0.27) as compared to Patescibacteria MAGs enriched in the 0.1 µm filter fractions (1.25 ± 0.029). When we compared the GRiD values of individual classes of Patescibacteria between 0.1 and 0.2 µm filter fractions, only MAGs from class Paceibacteria showed significantly higher GRiD values in the 0.2 µm filter fractions (Welch Two Sample t-test, p = 6.14e-03, Figure 5, B).

### Limited metabolic and biosynthetic capabilities in Patescibacteria

Metabolic reconstructions based on KEGG modules revealed that the metabolic repertoire of the analyzed Patescibacteria genomes did not show a clear separation by their taxonomy (Figure 6) nor followed a particular pattern in oxic and anoxic wells (Additional file 5, Figure S2). All Patescibacteria MAGs lacked central energy metabolism and biosynthetic pathways for most amino acids and vitamins. The tri-carboxylic acid (TCA) cycle was missing in 81.8% of the Patescibacteria MAGs and was incomplete for the remaining 18.2% of the MAGs. Glycolysis was incomplete in all MAGs, pentose phosphate pathway (PPP) was incomplete in 92% of the MAGs, and reductive PPP was absent in 97% of the MAGs. Biosynthesis pathways for most of the amino acids (except serine, glycine and sometimes asparagine) and vitamins (except cobalamin and thiamin) were missing in most of the Patescibacteria MAGs. In addition, electron transport chain complexes (I-IV) were not identified, with exception of gene encoding for the F- Type ATPase (from ETC complex V) in 59.7% of the Patescibacteria.

**Figure 6:**
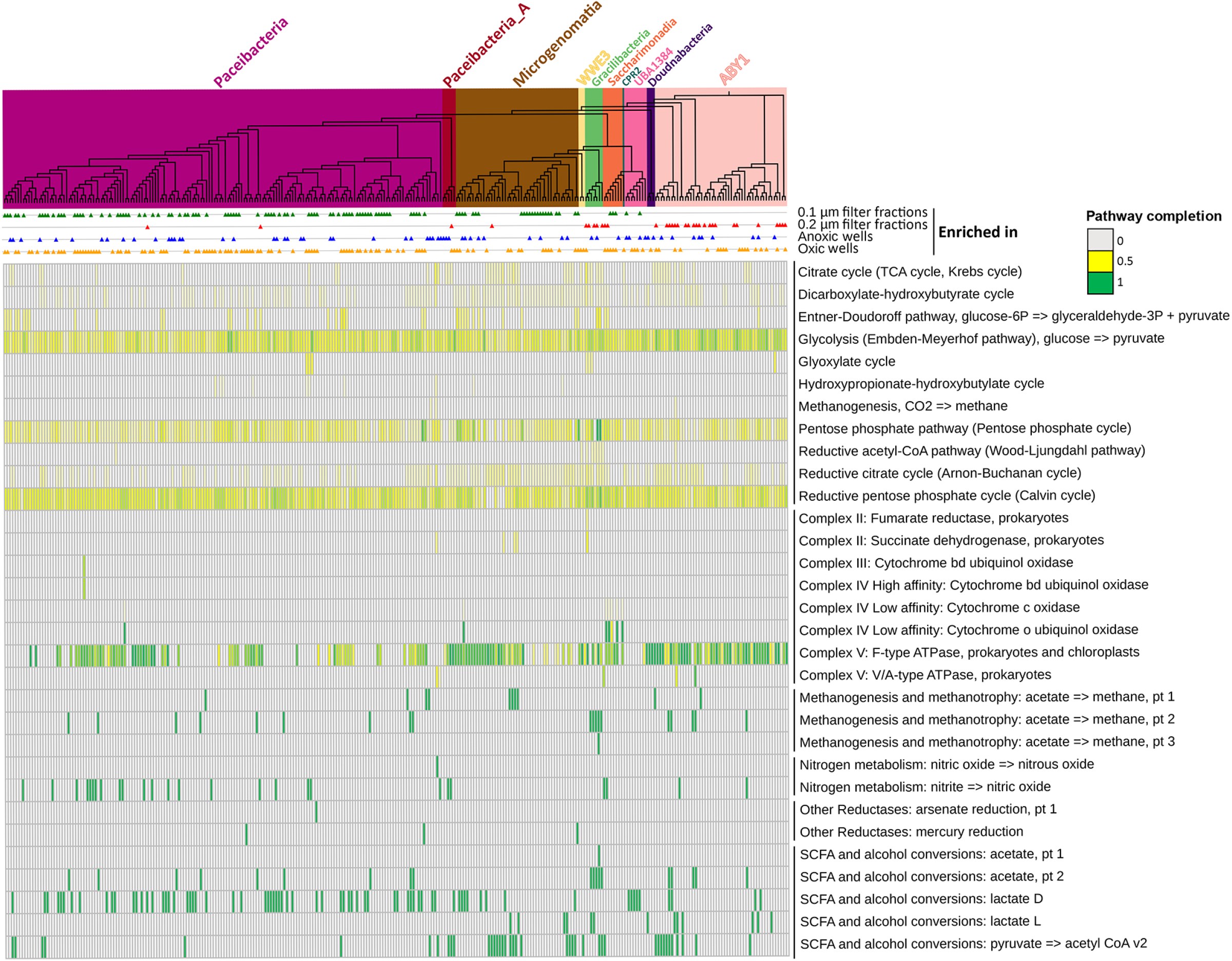
Metabolic and functional repertoire of the high quality Patescibacteria MAGs. The heatmap shows completeness of pathways and presence/absence of the functions in 291 high-quality Patescibacteria genomes annotated within DRAM [65], arranged according to their phylogenetic placement. Clade background colors within the phylogenetic tree represent respective taxonomic classes of Patescibacteria. Colored triangles next to each genome represent their enrichment in 0.1 µm filter fractions (green), 0.2 µm filter fractions (red), anoxic wells (blue) and oxic wells (orange), respectively. Electron transport chain complexes I-IV, sulfur metabolism functions, and photosynthesis related genes were absent from almost all the MAGs. A similar heatmap arranged as per the 5-fold enrichment of the MAGs in oxic and anoxic wells is provided as Additional file 5, Figure S2.

However, Patescibacteria possessed some notable genes, namely those coding for copper transporter (*copA*) and cobalt transporter (*corA*) that are usually found in pathogenic bacteria [38,39]. Also, carbohydrate active enzymes (CAZy) responsible for degradation of starch (11% MAGs), polyphenolics (25% MAGs) and chitin (11% MAGs) were observed. At least 13% of the MAGs had more than one type of CAZy. Patescibacteria also encoded genes for small chain fatty acids (SCFA) and alcohol conversion functions e.g. D-lactate dehydrogenase (25% MAGs), L-lactate dehydrogenase (4% MAGs), and conversion of pyruvate to Acetyl-CoA (K00174, 14% MAGs). Acetate kinase was found in only 6% of the Patescibacteria MAGs. A mutually exclusive presence of D- and L-lactate dehydrogenases was observed.

### Genomic signs of adaptive response of Patescibacteria to oxic and anoxic conditions

We classified 134 Patescibacteria MAGs as 5-fold enriched in oxic wells (H32, H41 and H51) and 64 Patescibacteria MAGs as 5-fold enriched in anoxic wells (H14, H43 and H52). No taxonomic preference for oxic or anoxic conditions was observed. Patescibacteria MAGs enriched in oxic sites showed some unique features with respect to their ability to resist oxidative stress. We found that superoxide dismutase genes (at least one of the *sodA*, *sodB*, *sodC*, *sodF*, *sodM*, *sodN* or *chrC* genes) were encoded by significantly higher proportion (82.8%) of the Patescibacteria MAGs enriched in oxic wells than in anoxic wells (65.6%) (two-proportions z- test, p = 8.8e-03), but there was no evidence for other stress regulator genes (*oxyR*, *soxR*, *soxS*, *rpoS*). There were no relevant metabolic pathways or genes specific to the 64 Patescibacteria MAGs enriched in anoxic wells (Additional file 5, Figure S2).

Correlation of the genomic coverages (relative abundances) of the Patescibacteria MAGs enriched in oxic wells with the dissolved oxygen concentrations revealed highly significant positive correlations for 28 MAGs (Additional file 6). Most of these MAGs belonged to class *Cand*. Paceibacteria (family UBA1539/*Yonathbacteraceae*) and genus GWC2-37-13 from order UBA1406/*Roizmanbacterales*. Most of these MAGs (82%) carried superoxide dismutase gene (K04564) essential for protection against free superoxide radicals in oxic environments.

We chose the most abundant, high quality Patescibacteria MAGs from oxic well H41 (H41- bin288, 0.1 µm filter fraction, relative abundance = 0.75% ± 0.15) and anoxic well H52 (H52- bin095, 0.1 µm filter fraction, relative abundance = 2.28% ± 0.37) as model organisms to illustrate the commonalities and divergences in their genomes (Figure 7). We also included the second most abundant Patescibacteria MAG from the same oxic well H41 (H41-bin049, 0.1 µm filter fraction, relative abundance = 0.41% ± 0.02) from the same taxonomic family as the anoxic representative. This was done to rule out the genomic differences due to the relatively distant evolutionary history of the first pair (H41-bin288 and H52-bin095). The representative MAGs H41-bin288 and H41-bin049 from the oxic well H41 showed positive correlations with oxygen (R = 0.88, p = 2.0e-02 and R = 0.75, n.s., respectively), while the representative MAG from anoxic well (H52-bin095) showed a negative correlation (R = −0.43, n.s.).

**Figure 7:**
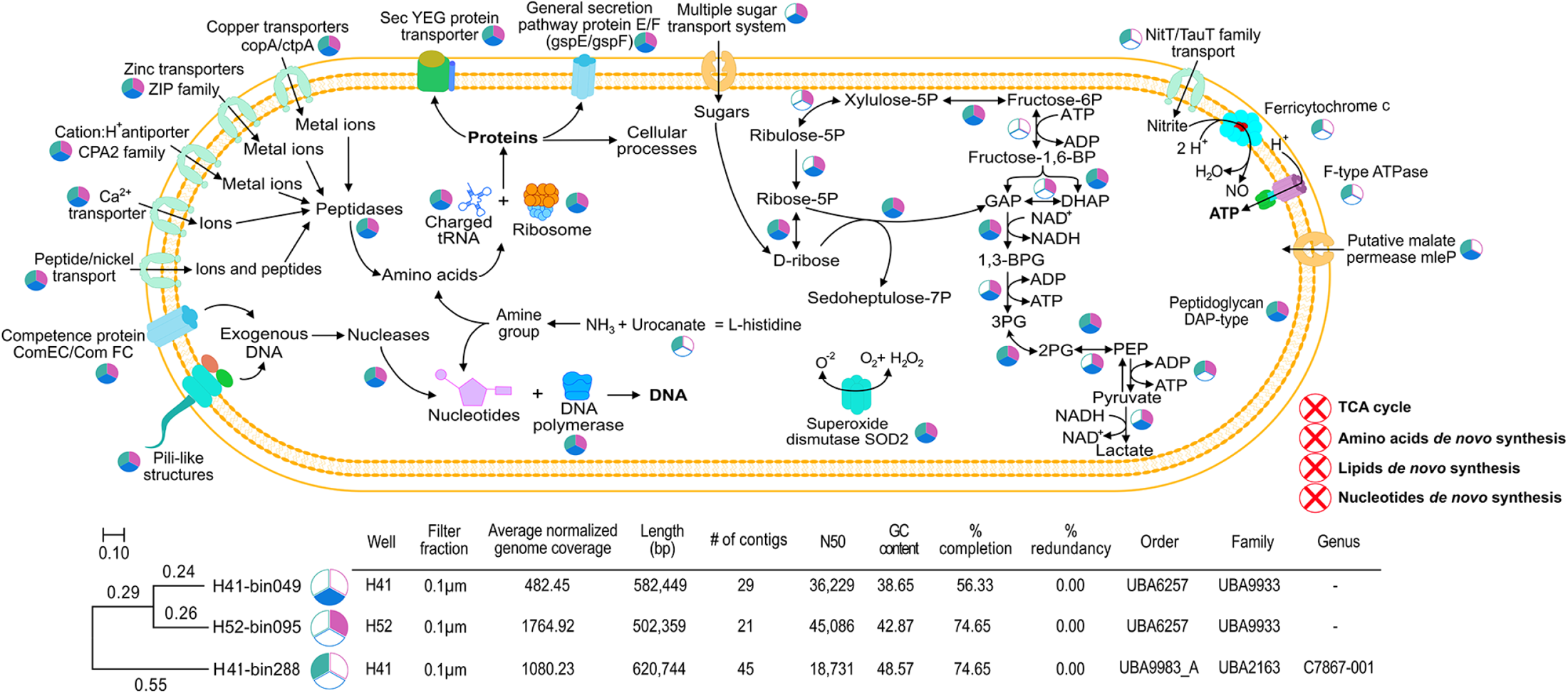
Cell schematic representing the functional repertoire of most abundant model Patescibacteria from oxic and anoxic groundwater wells. The common and genome specific gene features are shown for the three representative genomes based on KEGG pathways. The pie diagrams next to each reaction or function state the presence of respective enzymes or proteins in the three model organisms as per the color key (oxic representatives in green and blue, and the anoxic representative in pink), while absence is indicated by white color.

Features specific to both representative genomes from oxic well H41 were genes coding for F-type H^+^-transporting ATPase (subunit a, b, c, α, β and γ), NitT/TauT family transporter (involved in transport of inorganic ions like nitrate, sulfonate, and bicarbonate), and nitrite reductase (*nirK* involved in conversion of nitrite to nitric oxide). On the other hand, genes related to sugar sensing and multiple sugar transport systems (ABC.MS.S), and lactate dehydrogenase (fermentation) were specific to the anoxic representative. Common genes or functions were found for all three representative genomes, e.g. genes encoding type IV pilus assembly proteins (PilB, PilC, PilM, and PilO) as well as competence proteins (ComEC, ComFC), useful for DNA uptake from exogenous sources, superoxide dismutase (SOD2) for protection against superoxide radicals, transporters of metal ions like zinc, copper, calcium, nickel. We also identified genes encoding for rod-shape determining proteins, like RodA with additionally related genes encoding for proteins like MreB and MreC in the anoxic representative.

### Co-occurrence patterns of Patescibacteria with other microbial species

A co-occurrence network generated using metagenomic abundances of MAGs revealed that many species of Patescibacteria were consistently co-occurring with one another, as well as with species of other bacteria and archaea (Figure 8). The average normalized genome coverages for all the studied MAGs across both filter fractions of all the wells are provided in Additional file 7. The most common one-to-one associations were observed with MAGs from the phyla Nanoarchaeota (mostly order Pacearchaeales), Bacteroidota, MBNT15, and Bdellovibrionota. A small isolated cluster within the network showed indirect but close associations of Patescibacteria with multiple members of the phylum Nitrospirota (genus RGB.16.64.22), and phylum Omnitrophota (Figure 8).

**Figure 8:**
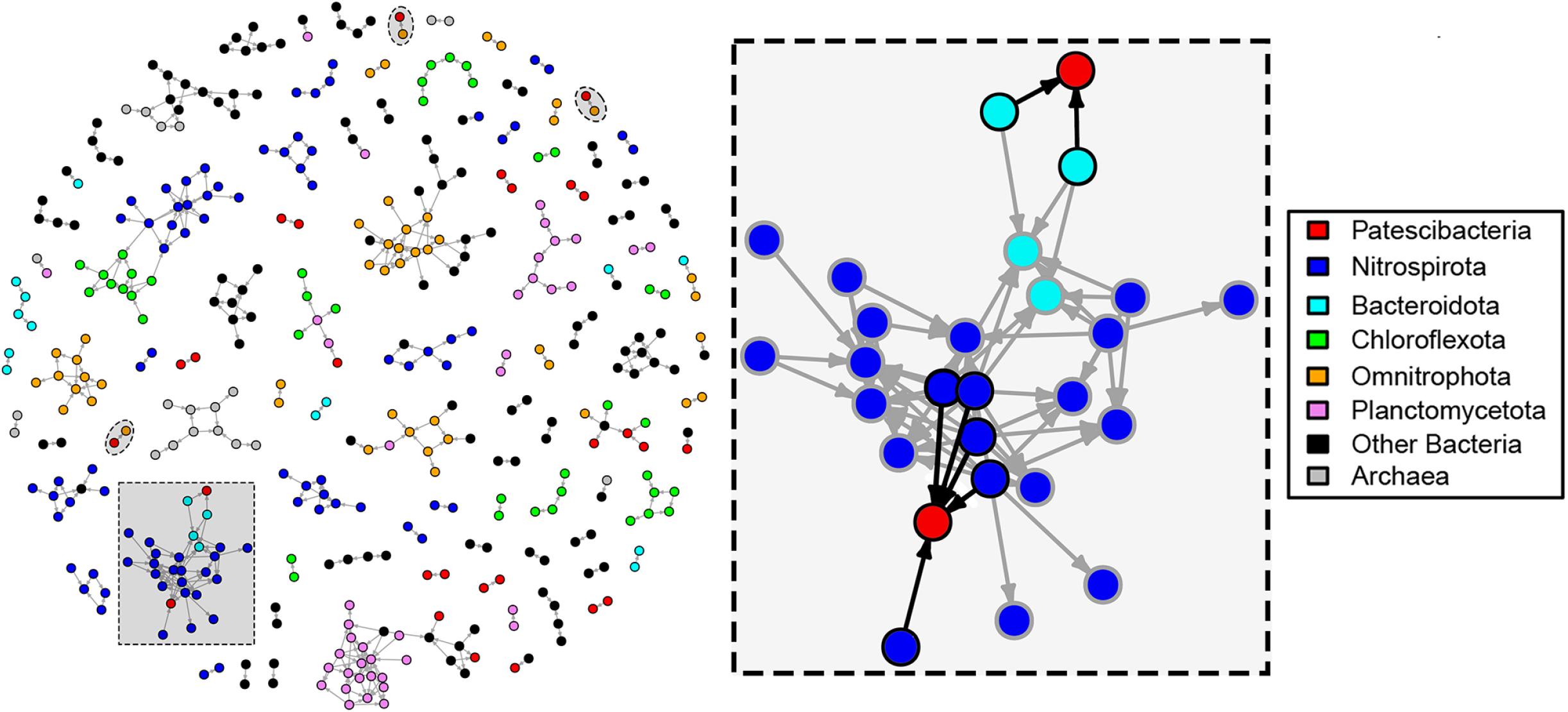
Co-occurrence network among the MAGs recovered from the studied groundwater wells. The proportionality network was constructed using normalized average coverages of the MAGs enriched (by 5-fold coverage difference) in 0.2 µm filter fractions as compared to 0.1 µm filter fractions to retain Patescibacteria possibly attached to other microbial hosts. The filled oval regions highlight the direct one-to-one associations of Patescibacteria MAGs paired with Omnitrophota MAGs. The zoomed-in cluster shows direct associations of Patescibcteria MAGs (filled red circles) with multiple Nitrospirota (filled blue circles) and Bacteroidota (filled cyan circles) MAGs highlighted with black outlines and arrows, while grey outlines and arrows indicate indirect associations. For construction of the proportionality network, ⍴ (rho) value cut-off of 0.95 was used.

Under the assumption that Patescibacteria were physically associated with larger host cells, we simplified our co-occurrence network to further refine the associations in the 0.2 µm filter fractions (using the 5-fold coverage cut-off as compared to 0.1 µm filter fractions). This follow- up co-occurrence network showed one-to-one associations of MAGs of the phylum Omnitrophota (class koll11) with MAGs from Patescibacteria (each one from the classes Paceibacteria, Microgenomatia, and candidate division ABY1). One of the MAGs from class Paceibacteria showed association with a Proteobacteria MAG (order Rickettsiales), while a MAG from candidate division ABY1 showed direct connections with two Bacteroidota MAGs. Another MAG from class Gracilibacteria showed direct connections with 5 Nitrospirota MAGs from the same genus UBA1546 (Figure 8). The sequence coverages of these highlighted genome pairs or clusters across the metagenomes are compared in Additional file 8, Figure S3 and Additional file 9, Figure S4. Two Actinobacteria MAGs belonging to the species *Aurantimicrobium* sp003194085 also showed associations with Patescibacteria. The first *Aurantimicrobium* interacted with a Patescibacteria (*Cand*. Paceibacteria) MAG, and the second with multiple Patescibacteria (2 *Cand*. Paceibacteria, 2 *Cand*. Gracilibacteria and 3 candidate division ABY1) MAGs.

When we searched for sequence similarity of all gene open reading frames (ORFs) from all Patescibacteria MAGs to ORFs from all other bacterial and archaeal MAGs in the present study using blastn [40], we found various ORFs from other taxa highly similar to Patescibacteria ORFs (95% sequence identity covering 85% length of the query and hit sequences). The most ORFs that matched were between members of genus UBA10092 of Patescibacteria (class Paceibacteria) and two members of the family UBA12090 of Omnitrophota (34 and 16 ORFs, respectively). They included genes encoding for twitching motility protein PilT (K02669), P- type Cu+ transporter (K17686) and lipopolysaccharide export system permease protein (K11720). Between members of genus UBA11707 of Patescibacteria (class ABY1) and genus UBA1573 of Proteobacteria (family Micavibrionaceae), 14 such ORFs, including gene encoding for ABC-2 type transport system ATP-binding protein (K01990), were observed. Thirteen such ORFs, including gene for ABC-2 type transport system permease protein (K01992), were observed between members of the family Zambryskibacteraceae of class Paceibacteria and genus ASMP01 of Nanoarchaeota.

To have an idea about the temporal co-occurrence patterns of other groundwater microbes with Patescibacteria, we additionally utilized time-series data based on 16S rRNA gene amplicon sequencing from the same groundwater transect from three wells (H41, H43 and H52) measured over more than six years [41]. We observed that Patescibacteria co-occurred mostly with members of phyla Proteobacteria (mostly order Burkholderiales) and Nitrospirota (order Thermodesulfovibrionia), in the well H41; Verrucomicrobiota, in the well H43 and Planctomycetota (mostly genus *Brocadia*) in the well H52. Similarly, a Patescibacteria MAG was identified to co-occur with multiple Thermodesulfovibrionia MAGs belonging to the phylum Nitrospirota in this study.

## Discussion

Our comprehensive metagenomic analyses revealed that modern pristine groundwater of the Hainich CZE is clearly dominated by *Cand*. Patescibacteria with an average relative abundance of 50% across all wells and a maximum of 79% in the 0.1 µm filter fraction. Compared to other groundwater communities dominated by CPR bacteria ranging from 2-28% [16], 3-40% [18], 10- 28% [7] and 36-65% [15], the exceptionally high abundance of CPR bacteria discovered in this study is distributed over distinct geochemical zones spanning oxic and anoxic conditions [17,33]. Although the spatial distribution patterns of the different *Cand*. Patescibacteria taxa (Figure 2) were less pronounced than those observed in other bacteria in groundwater of the Hainich CZE [33,41], and despite their streamlined genomes, we could highlight certain environmental preferences of the *Cand*. Patescibacteria. Access to 587 manually curated MAGs of *Cand*. Patescibacteria, assigned to different filter fractions, allowed us to shed some light on genomic characteristics linked to their cell size and a putative free living or host attached lifestyle.

Patescibacteria have been described mostly in anoxic or hypoxic environments [42,43]. Our data show no major metabolic or taxonomic differences in Patescibacteria enriched in oxic and anoxic groundwater wells. Significantly higher proportion of superoxide dismutase genes in Patescibacteria MAGs enriched in oxic groundwater wells compared to those in anoxic wells is an example of spatial differentiation that might be due to an environmental selection mechanism, as these enriched species have an advantage to withstand the presence of oxygen radicals when exposed to high O2 concentrations. More than 80% of the Patescibacteria MAGs enriched in oxic wells could potentially resist superoxide radicals, and more than 20% showed a positive correlation to oxygen concentrations, in particular those belonging to class *Cand*. Paceibacteria (family UBA1539/*Yonathbacteraceae*) and to order UBA1406/*Roizmanbacterales*. But even closely related Patescibacteria species showed different preferences for oxygen concentrations in terms of metabolic pathways (Figure 7).

The permanently high O2 concentration in well H32 (2.23 ± 0.56 mg/L) and especially in well H41 (4.83 ± 1.7 mg/L) [41,44], did not lead to enrichment of groundwater Patescibacteria MAGs with genetic traits of energy harvesting mechanisms through aerobic respiration. Exposure to oxygen is not exceptional for *Cand.* Patescibacteria, as oxic soils are the main source for their vertical translocation into shallow groundwater [17,24]. *Cand.* Patescibacteria represent only 0.55% of the total bacterial soil community in the preferential forest surface-recharge area of the Hainich CZE (Herrmann *et al.* 2021, unpublished observations). Despite this low abundance, these ultra-small organisms are readily mobilized from soil, especially during winter months when ionic strength of the seepage is very low (Herrmann *et al*. 2021, unpublished observations), and as such constitute the largest fraction of taxa shared between seepage and shallow groundwater [17].

The most abundant Patescibacteria MAG from oxic well H41 (H41-bin288) had genes that encode for nitrite transport and its subsequent reduction into nitric oxide involving ferricytochrome c. Also, this genome possessed a gene for F-Type ATPase to generate energy by ATP formation and it did not encode genes for fermentation (L- or D-lactate dehydrogenase).

This collectively suggests the possibility of an alternative anaerobic respiration mechanism in this particular genome. Despite the low *in situ* concentrations of nitrite, it might be alternatively provided by the nitrification process. This relates to the fact that well H41 is characterized as a nitrification hotspot with measured rates of 0.48 ± 0.09 and 0.64 ± 0.39 nmol NOx liter^−1^ h^−1^ [45] and to the high relative abundances of *Nitrospira* on the metagenome level and *Thaumarchaeota* on the metatranscriptome level [46]. Presence of genes coding for multiple subunits of F-Type (H^+^ transporting) ATPase in this genome confirms the existence of supplementary ATP synthesis machinery, which are commonly observed in aerobic bacteria [47]. Similarly, notable features specific to both representative genomes from oxic well H41 included genes involved in the transport of inorganic ions like nitrate, sulfonate, and bicarbonate.

The almost complete absence of the aerobic respiration machinery i.e. the electron transport chain complexes, terminal oxidases / electron acceptors, and gene products associated with the TCA cycle, along with widespread presence of L- or D-lactate dehydrogenases confirms the previously postulated fermentative lifestyles of Patescibacteria [11,15,48] in members of the three lineages OD1 (Parcubacteria), OP11 (Microgenomates), and BD1-5 (Gracilibacteria). Parcubacteria were proposed to produce acetate, ethanol, lactate, and hydrogen as fermentation products based on metagenomic and proteomic analysis [3,15,48]. Presence of L- or D-lactate dehydrogenase genes in one third of the Patescibacteria MAGs indicates specificity for fermentation substrates. In one tenth of the MAGs enriched in 0.1 µm filter fractions, specificity for L-lactate could be observed based on the exclusive presence of L-lactate dehydrogenase genes. Presence of multiple carbohydrate active enzymes (CAZy) in many Patescibacteria suggests their potential for degradation of multiple complex compounds like starch, chitin, and polyphenolics.

The spatial differentiation of *Cand.* Patescibacteria could also be indirectly caused by the preference of a putative host organism for certain environmental conditions. The oxic, nitrate- rich (15.71 mg/L) groundwater of well H41 was dominated by Nitrospirota MAGs, and 5 of them co-occurred with a single Patescibacteria MAG (H52-bin081_1, *Cand.* Gracilibacteria) and had similar abundance patterns (Additional file 9, Figure S4). As some Nitrospirota MAGs (n = 51) were enriched exclusively in oxic wells, their preference might have determined the distribution pattern of putative CPR episymbionts. Nitrospirota species were also found to be consistently co-occurring with Patescibacteria in some of the studied wells based on OTU abundances from 16S rRNA gene amplicon sequencing data collected over 6.5 years [41] as well as MAG abundances from this study across the groundwater transect. At the minimum, these observations suggest common niche preferences between some members of these two phyla.

To elucidate other possible associations of Patescibacteria with other prokaryotes, we utilized above mentioned time-series data that revealed consistent co-occurrence of Patescibacteria OTUs with OTUs from Proteobacteria, Verrucomicrobiota, and Planctomycetota in addition to OTUs from Nitrospirota [41]. When we looked into the genomic characteristics of all Patescibacteria and all other MAGs, we found various ORFs from other taxa highly similar with Patescibacteria, between members of (i) class Paceibacteria and family Omnitrophota, (ii) class ABY1 and family Micavibrionaceae, and (iii) family Zambryskibacteraceae of class Paceibacteria and genus ASMP01 of Nanoarchaeota, suggesting probable acquisition of motility and transport functions from other bacteria or archaea.

Network analysis based on abundances of all MAGs of both filter fractions revealed that the members of the phyla Bacteroidota, MBNT15, and Bdellovibrionota along with members of phyla Nitrospirota and Omnitrophota had direct specific connections with some Patescibacteria. Furthermore, we restricted the network analysis only to MAGs enriched on the 0.2 µm filter fractions (57 Patescibacteria and 423 other MAGs) in order to identify Patescibacteria that would be potentially attached to other larger host cells. This narrowed-down analysis showed interactions of Patescibacteria with few specific MAGs of the phyla Bacteroidota, Nitrospirota, Omnitrophota, and Actinobacteria. Our co-occurrence analysis did not reveal direct connections of Actinobacteria MAGs with any of the Saccharibacteria, although Actinobacteria are reported as host for Saccharibacteria (TM7) in human oral cavity [12,27,29]. However, direct network connections of *Aurantimicrobium* species, members of the phylum Actinobacteria with multiple other Patescibacteria MAGs from classes Paceibacteria, Gracilibacteria, and candidate division ABY1 hint towards possible host-symbiont relationships in these particular pairs.

Direct one-to-one connections with members of other phyla were found in only 5 out of 57 (8.77%) Patescibacteria MAGs enriched in 0.2 µm filter fractions, suggesting that the majority of groundwater Patescibacteria of the Hainich CZE is not specifically associated with one single host, but associations with multiple hosts cannot be ruled out. The attachments between cells are often fragile and may be partly or completely disrupted during filtration and sample processing steps, and hence are difficult to track using sequential filtration. An even lower percentage of associations (<1.5%) based on potentially co-sorted SAGs containing DNA from heterogeneous sources was reported from Beam *et al*. 2020 [11].

On average, Patescibacteria enriched in 0.1 µm filter fractions had 22% smaller genome size than those enriched in 0.2 µm filter fractions, and it has been previously shown that smaller cell size is linked to genome reduction [49,50]. This genome size difference might be due to differences in average cell sizes of *Cand*. Paceibacteria and *Cand*. Microgenomatia that were preferentially enriched within 0.1 µm filter fractions; and candidate division ABY1, and *Cand*. Gracilibacteria that were preferentially enriched within the 0.2 µm filter fractions. Smaller genomes in tiny CPRs might be the result of genome streamlining leading to lack of complex energy metabolism and biosynthetic capabilities which makes them rely on other cells through cell-cell attachment.

We found Type IV pilus assembly proteins in a higher proportion of Patescibacteria enriched in 0.1 µm filter fractions. These proteins are responsible for formation of pilin-like appendages that are involved in a variety of functions like adherence to host cells, locomotion, DNA uptake as well as protein secretion in bacteria [51], which would support physical association with other microbes. Type IV pili (T4P) are essential for virulence of some Gram-negative pathogenic bacteria [52] and also found in Gram-positive bacteria with a different pilus assembly mechanism involving a sortase [53]. Pili like appendages were microscopically shown to form surface attachment of CPR bacteria with other (host) large cells [18]. The symbiotic association of TM7i (*Cand*. Saccharibacteria) with its host *Leucobacter aridocollis* J1, mediated by T4P was identified in a co-culture experiment [54]. As pilus mediated attachments are often fragile, small Patescibacteria cells passing through the 0.2 µm filters do not necessarily indicate lack of cell-cell attachment with larger bacterial cells. Many of these ultra-small Patescibacteria appear to have a rod-shaped morphology, as genes encoding the rod shape-determining protein (MreB) were found in a higher proportion of MAGs enriched in 0.1 µm filter fractions. The recent reconstruction of the last bacterial common ancestor (LBCA) genome of CPR lineage suggests a rod-shaped morphology [55]. However, most of the reported morphologies for the Patescibacteria are cocci [12,18,21]. Although we cannot rule out that some of the larger rod-shaped Patescibacteria could still pass through the 0.2 µm filter pores, this would not explain the enrichment in the 0.1 µm filter fractions. More direct microscopic visualization is needed to verify the morphology of these ultra-small Patescibacteria.

We found higher growth rates of Patescibacteria in near-surface wells (H14, H32) of the groundwater transect than in the ones more downstream. Growth of CPR bacteria is stimulated after attachment to host-cells [18]. As cell-cell aggregations might be more prone to dispersal limitations in a dense rock matrix, surface-near wells could have higher probabilities of host interactions. But our co-occurrence analysis did not reveal direct connections of CPR MAGs with higher growth rates with other MAGs.

Groundwater of the very shallow well H14, located uphill of the transect, shows a fast response to weather events [56], and is characterized by both the highest bacterial diversity and the presence of well-known surface heterotrophs; whereas core groundwater species dominated groundwater microbiomes in the downstream direction [33]. This well, along with the other near-surface well (H32) showed the lowest relative abundances of Patescibacteria and of Patescibacteria MAGs, although those that were detected had higher expected replication rates on average. A possible explanation for this pattern is that surface exported members were replicating within the soil before being flushed into the groundwater. Other, more successful groundwater CPR groups may have slower growth and replication rates within the transect due to much lower microbial cell densities and less available organic carbon. Indeed, some taxa such as those belonging to *Cand.* Saccharimonadia, which had among the highest growth rates, did not flourish within other wells of the groundwater transect. We hypothesize that they might be more adapted to soil habitats, which was also observed in previous studies [17].

The predominance of particular CPR species in oxic (H41) and anoxic (H52) wells appears to be the result of environmental preference or exploitation of other organisms for cellular requirements in the nutrient deficient groundwater. Some potential hosts supporting an episymbiotic lifestyle could be identified. The environmental preference of some of these hosts, e.g. Nitrospirota for oxygen and nitrogen in well H41, would explain the predominance of their potential Patescibacteria episymbiont in H41, with an estimated episymbiont-to-host ratio of 3.6:1 based on coverages of Patescibacteria and Nitrospirota MAGs in total coverage of all binned genomes. But the vast majority of the ultra-small Patescibacteria in the groundwater appears to be free-living, self-sufficient with their minimal genomes [11,42], adapted to oligotrophic conditions with low growth rates, and equipped with genes to cope with oxidative stress only if needed. We found evidence that the majority has the capability to attach to other cells, which appears to also include other Patescibacteria, and this attachment might be not very specific or for longer time periods, just long enough to loot or exchange supplies.

## Conclusions

The Candidate Phyla Radiation represent the largest phylogenetic diversity within the bacterial domain, which has not been reflected in the metabolic versatility of genomic representatives studied to date. Here we leveraged a well characterized aquifer transect, that is dominated by members of the CPR and spans large biogeochemical gradients, to explicitly explore genomic adaptations to environmental conditions. The most significant and surprising result was the high level of similarity in predicted metabolic functions and expected lifestyles that spanned large redox gradients from fully oxic to completely anoxic groundwater, both within the larger CPR clade as well as at finer phylogenetic resolutions. One noteworthy exception was a differential abundance in superoxide dismutase, a potentially useful indicator of oxygen exposure in CPR genomes recovered from other environments or already deposited to sequence databases. Due to a suspected dependence on other bacterial hosts, we searched among >1200 constructed MAGs and a larger amplicon dataset for potential partners, finding that only 8% of CPR MAGs exhibited significant one-to-one relationships. Therefore, we propose that most members of the CPR form non-specific associations, attaching to multiple hosts to supplement their energetic demands within oligotrophic groundwaters.

## Methods and Materials

### Groundwater sampling, DNA extraction and sequencing

Samples were collected from a groundwater transect system spanning through a ∼6 km long zone including forest, pasture and agricultural land within the Hainich Critical Zone Exploratory (CZE) located in Thuringia, Germany. The Hainich CZE was established and extensively studied by Collaborative Research Center AquaDiva [32]. The groundwater was collected from 6 wells (H14, H41, H43, H51, H52) in January 2019 and (H32) in November 2018 spanning various zones of the transect. For each well, on average 61.3 ± 35.4 liters of groundwater was filtered through 0.2 µm filters (Omnipore Hydrophilic PTFE membrane, Merck Chemicals GmbH) followed by 0.1 µm filters in triplicates (except for well H32 where there were only two replicates out of which one from the November 2018 sampling campaign was used as biological replicate). All the 32 filter fractions were immediately frozen and stored under -80°C. The DNA was extracted using a phenol/chloroform protocol, the libraries generated with an NEBNext Ultra FS DNA preparation kit, and sequenced on an Illumina NextSeq 500 system with paired-end library (2 × 150 bp).

On an average 9.8 ± 1.15 Gb of raw DNA sequence data were obtained from each of the 32 filter fractions. Of which, 86.12 ± 0.57 % of the reads were of very high quality (at least quality score Q40). Subsequent quality control steps like adapter trimming, PhiX detection and removal using BBDuk (bbtools version 37.09, written by Brian Bushnell, last modified March 30, 2017) further improved the quality of the reads. These high-quality reads were then used for metagenomic assembly and followed by genome binning steps.

### Metagenomic assembly, genome binning and refinement

The quality controlled reads of each individual filter fraction replicate were assembled and scaffolded using metaSPAdes v3.13 [57]. Scaffolds larger than 1 kb were used for downstream analyses. Genome binning was carried out using three binning algorithms - Abawaca v1.07 [15], ESOM [58,59] and Maxbin2 v2.2.4 [60]. The values 3000 and 5000 bp as well as 5000 and 10000 bp were used as *-min* and *-max* parameters to calculate 4-mer frequencies for Abawaca and ESOM (the script esomWrapper.pl, https://github.com/tetramerFreqs/Binning), and both the 40 and 107 marker gene sets were utilized in Maxbin2. DASTool v1.1 [14] was used to determine the best bins among these approaches. Bins were further refined manually inside the Anvi’o workflow v6.1 [35,36]. The quality of the refined bins (completeness and contamination/redundancy) was also calculated based on domain-level single-copy core genes within Anvi’o. Genomes from each assembly were de-replicated using dRep v2.6.2 [61] at 99% ANI to remove strain level redundancy across sites, resulting into 1275 representative MAGs. Genome coverages were calculated within Anvi’o, and were normalized using number of RNA polymerase B (*rpoB*) genes identified within the metagenomic reads.

### Taxonomic assignments, gene annotations and pathway predictions

Overall community composition of each metagenome was determined using phyloFlash v3.4 [62] based on proportions of reads mapped to SILVA SSU rRNA Ref NR99 database, Release 138 [34]. Taxonomic classification of individual MAGs was performed by GTDB-Tk v0.3.2 [63] using GTDB Release 89 as reference database. Out of the 1275 genomes GTDB-Tk classified 587 genomes as *Cand*. Patescibacteria at phylum level. We used *anvi-script-gen-CPR-classifier* script from Anvi’o v6.1 [35,36] which uses supervised machine learning model (random forest classifier) to train the program and *anvi-script-predict-CPR-genomes* for predicting the probability of the MAGs to confirm the CPR genomes. The training is based on the profile of previously published 139 single copy core genes from hundreds of CPR genomes from Brown *et al.* [15] and Campbell *et al.* [37] as input. This model confirmed 291 out of 587 genomes as CPR with a high confidence score (75% or more). While the model was inconclusive in case of the remaining 174 genomes based on low confidence score (less than 75%) and the remaining 122 genomes were discarded due to their completion levels below 50%.

The gene annotations, coding sequences, respective protein sequences, coverage calculations and other mapping statistics for all the genomes were exported by *anvi-summerize* program from within the Anvi’o workflow. The annotations were also carried out using Prodigal v2.6.3 [64]. Distilled and Refined Annotation of Metabolism (DRAM) [65] was used to generate pathway / metabolism summaries. At least one proper (other than hypothetical, uncharacterized or gene with unknown function) annotation from KEGG [66], MEROPs [67], Pfam [68] or dbCAN [69] was considered. This generated a single tab delimited annotation file listing the best hits from all these databases as well as summaries focused on most important pathways and functions. The pathway coverages (completeness) of central metabolism pathways were calculated based on KEGG modules definitions (https://www.genome.jp/kegg/module.html).

### Phylogenetic analysis

Single copy core bacterial genes were detected in all the 1087 bacterial MAGs using hmm profile (default ‘Bacteria_71’ hmm profile in Anvi’o v6.1), their protein sequences were extracted and aligned using MUSCLE [70] from within the Anvi’o [35,36]. A phylogenetic tree based on multiple sequence alignment of the 68 core proteins present in all bacterial MAGs (1087) was constructed using Approximate Maximum Likelihood in FastTree v2.1.11 SSE3, OpenMP [71] with 1000 bootstrap replications. The subset of the tree was used for arranging the metabolic pathways of 291 selected Patescibacteria MAGs in Figure 6.

### In-situ measurement of replication

The forward sequencing reads from all the metagenomes were mapped to the MAGs to calculate the sequence coverage of individual contigs. These coverage profiles were utilized to calculate Growth Rate InDex (GRiD) [72] which is directly proportional to the growth rates of the cells in a given environment. GRiD measures the difference in genome copies closer to the origin of replication compared to the terminus caused by ongoing replication forks. The coverage cut-off of 0.7 was used to remove extremely low coverage contigs.

### Statistical analyses

The difference in the mean genome sizes of the MAGs enriched in different filter fractions were compared using Kruskal-Wallis rank sum test followed by pairwise Dunn’s test in R [73]. The proportions of gene annotations (KEGG) in the MAGs enriched in different filter fractions or oxic and anoxic wells were compared with two-proportions z-test with Yates’ continuity correction in R. The p-values were adjusted for multiple testing using ‘fdr’ correction unless otherwise mentioned.

### Co-occurrence network analysis

We used normalized average genome coverages of all the 1275 MAGs across all the metagenomes as the approximation of abundance profiles of species from respective metagenomes. This abundance matrix was used to calculate proportionality of the coverage profiles in R package propR v4.2.6 [74]. A *⍴* cutoff of 0.95 was used for network creation to highlight only the most relevant co-occurrences. The network was generated using the R package igraph v1.2.6 [75] and exported to Cytoscape v3.8.2 [76] for visualization using R package Rcy3 v2.8.1 [77].

### Search for ORF similarity

We carried out blastn [40] search on all the annotated ORFs for Patescibacteria MAGs as a query against all the ORFs of all the MAGs other than Patescibacteria. We filtered the results based on 95% sequence identity over 95% query and hit ORF length with e-value cut off of 1.0e-5. We chose only one hit in case of more than one hits for the same query sequence.

## Declarations

### Availability of data and material

Data used for this study were deposited into the European Nucleotide Archive (ENA). The raw metagenomic sequencing reads were deposited under ENA project accession PRJEB36505, assemblies for individual samples were deposited under ENA project accession PRJEB36523.

### Authors’ contributions

NMC, KK, WAO, MT, AJP designed this study. WAO, KK, AJP, TLVB, and MT planned, designed, and conducted the metagenomic sampling approach. MM, MH helped during metagenomic sequencing. NMC, WAO, TLVB, and AJP performed the metagenomic analysis. NMC manually curated and performed comparative genome analysis of the MAGs. PAFG conducted the metabolic reconstruction analysis of representative MAGs. NMC, KK, WAO wrote the manuscript with the help of all authors.

## Acknowledgements

We thank Patricia Geesink and Falko Gutmann for filtration and DNA extraction of groundwater samples, and the Hainich CZE site manager Robert Lehmann for their assistance with sample preparation, collection, and filtration. Additionally, we thank Ivonne Görlich and Marco Groth from the Core Facility DNA sequencing of the Leibniz Institute on Aging - Fritz Lipmann Institute in Jena for their help with Illumina sequencing. We also thank Syrie Hermans for providing preliminary data from time-series analysis of 16S rRNA gene amplicon sequencing for some of the groundwater wells.

## Additional files

**Additional file 1:** Single copy genes from publically available CPR genomes used to predict CPR MAGs in this study. The file was taken from Anvi’o codebase (https://github.com/merenlab/anvio).

**Additional file 2:** Genome statistics and taxonomic assignments of Patescibacteria MAGs in this study.

**Additional file 3:** The Newick tree file for phylogenetic tree shown in Figure 3, B.

**Additional file 4, Figure S1:** Correlation of average normalized genome coverages of Patescibacteria MAGs from respective wells with respective GRiD values.

**Additional file 5, Figure S2:** Metabolic and functional repertoire of high quality Patescibacteria MAGs. The heatmap shows completeness of pathways and presence/absence of functions in 291 high-quality Patescibacteria genomes annotated with DRAM, arranged according to their enrichment in oxic and anoxic wells based on 5-fold coverage criterion.

**Additional file 6:** Correlations of average normalized genome coverages of Patescibacteria MAGs enriched in oxic wells with dissolved oxygen and nitrate concentration.

**Additional file 7:** Genomic coverages of 1275 microbial MAGs in all studied metagenomes.

**Additional file 8, Figure S3:** Coverage distribution of selected MAGs from the network in Figure 8. Only the direct one-to-one pairs of Patescibacteria with other MAGs are plotted.

**Additional file 9, Figure S4:** Coverage distribution of selected MAGs from the highlighted cluster in network in Figure 8. Only the direct connections of Patescibacteria with other MAGs are plotted.

