## Supplementary figures and images for "The economical lifestyle of CPR bacteria in groundwater allows little preference for environmental drivers"

### Additional file 4, Figure S1

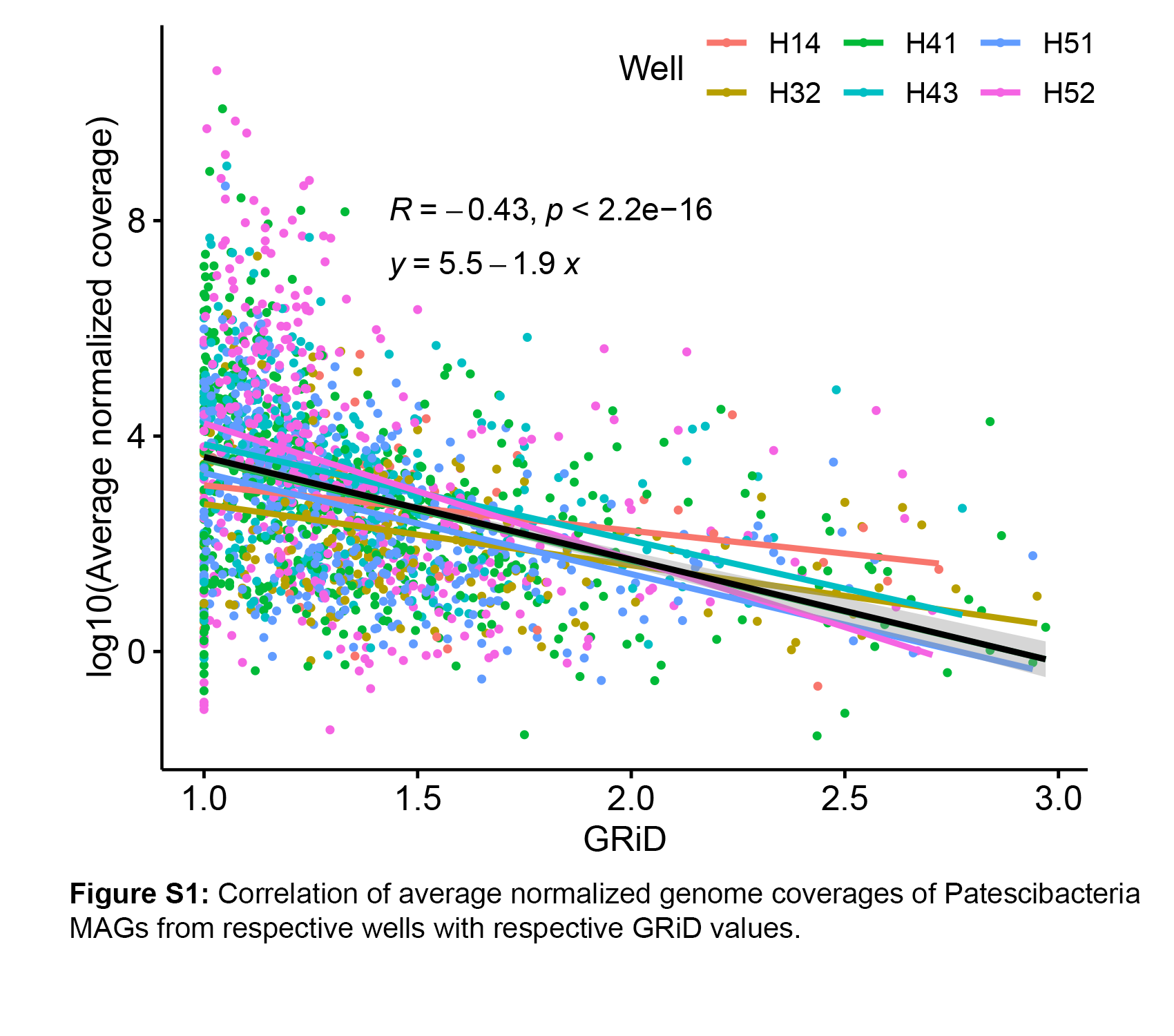

### Additional file 5, Figure S2

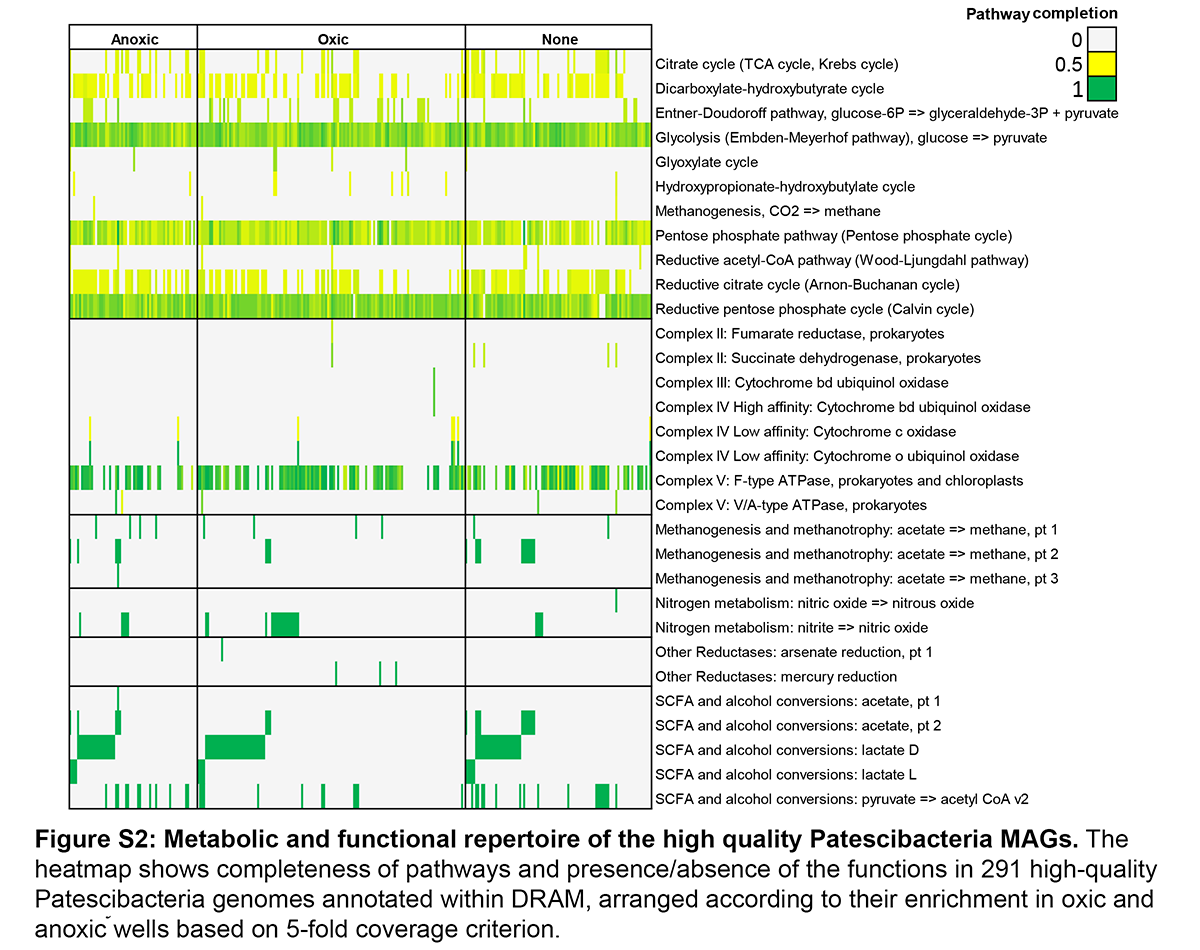

### Additional file 8, Figure S3

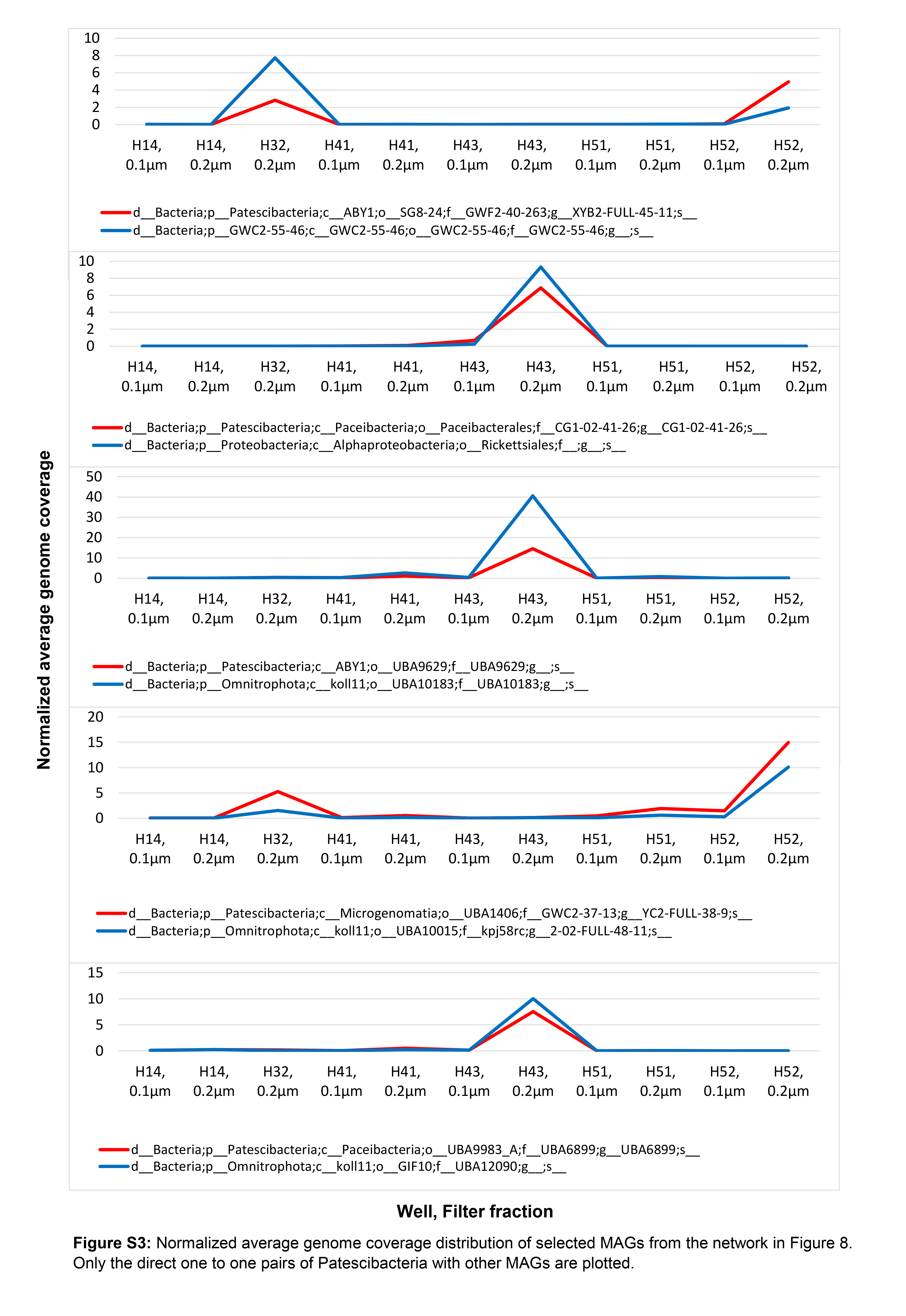

### Additional file 9, Figure S4

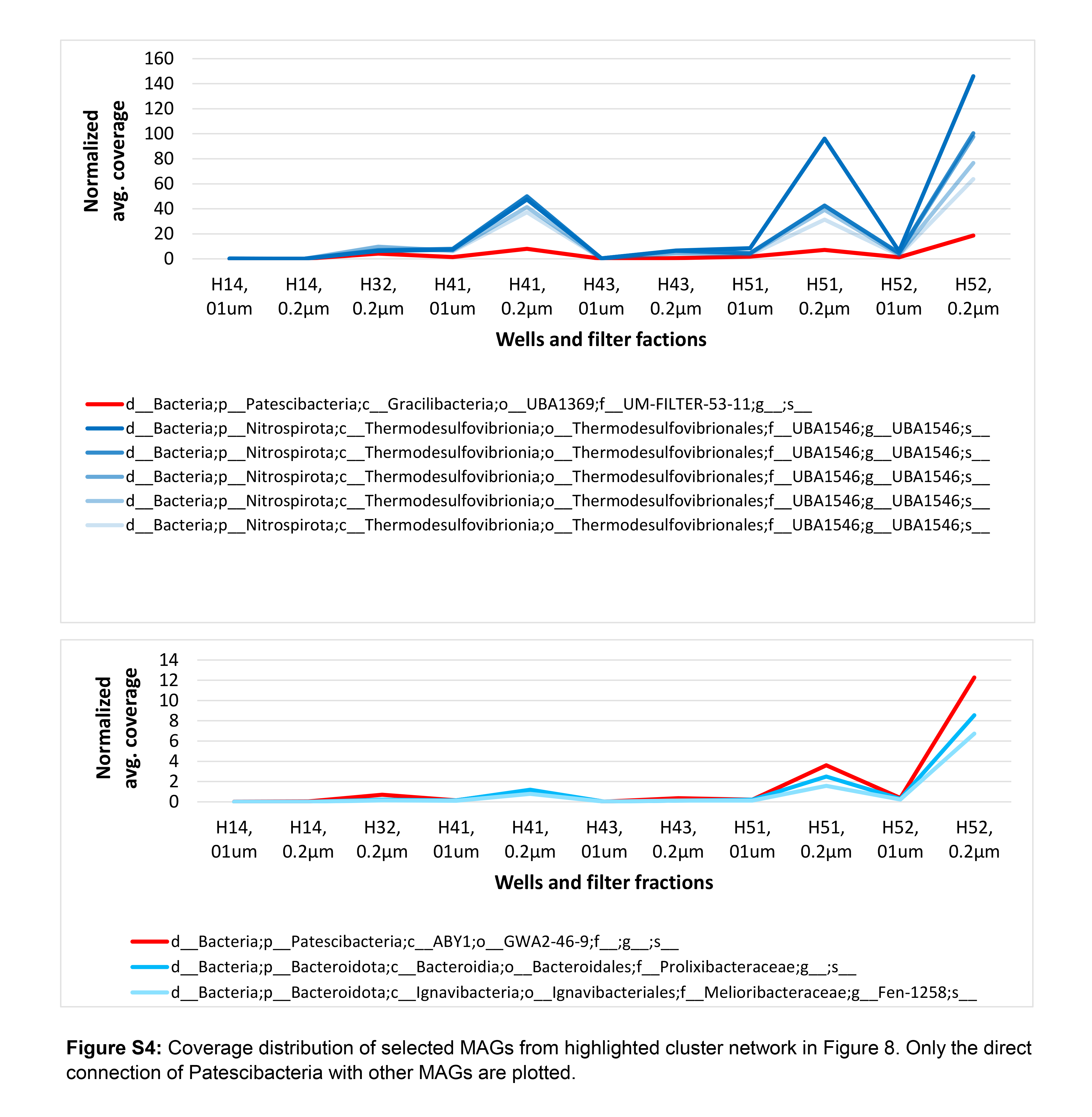
